# AML ImmunoPET: ^64^Cu-DOTA-anti-CD33 PET-CT imaging of AML in Xenograft mouse model

**DOI:** 10.1101/635078

**Authors:** Srideshikan Sargur Madabushi, Jamison Brooks, Darren Zuro, Bijender Kumar, James Sanchez, Liliana Echavarria Parra, Marvin Orellana, Paresh Viswasrao, Indu Nair, Junie Chea, Kofi Poku, Nicole Bowles, Aaron Miller, Todd Ebner, Justin Molnar, Joseph Rosenthal, Daniel A Vallera, Jeffrey Y C Wong, Anthony S Stein, David Colcher, John E Shively, Paul J Yazaki, Susanta K Hui

## Abstract

**Purpose:** Acute myeloid leukemia (AML) is a highly aggressive form of leukemia that results in poor survival outcomes. Currently, diagnosis and prognosis are based on invasive single-point bone marrow biopsies (iliac crest). There is currently no AML-specific non-invasive imaging method to assess disease in the whole body, including in extramedullary organs, representing an unmet clinical need. About 85-90% of human myeloid leukemia cells express CD33 (an accepted biomarker for AML) cell surface receptors, highlighting CD33 as an ideal candidate for AML immunoPET.

**Experimental Design:** In this study, we evaluated if ^64^Cu-DOTA-anti-CD33 murine monoclonal antibody (mAb) can be used for ImmunoPET-CT imaging of AML in preclinical model. For translational purpose, a humanized anti-CD33 antibody was produced, and after confirming its anti-CD33 PET-CT imaging ability we further explored its therapeutic potential.

**Results:** ^64^Cu-DOTA-anti-CD33 based PET-CT imaging detected CD33^+^ AML in mice with high sensitivity (95.65%) and specificity (100%). In the entire skeletal system, CD33^+^ PET activity was significantly high in the bones of AML bearing mice over non-leukemic control mice; femur (p<0.00001), tibia (p=0.0001), humerus (p=0.0014), and lumber spine (p<0.00001). Interestingly, the imaging showed CD33^+^ PET activity prevelant in epiphysis/diaphysis of the bones, indicating spatial heterogeneity. Anti-CD33 therapy using newly developed humanized anti-CD33 mAb as an ADC (p=0.02) and ^225^Ac-anti-CD33-RIT (p<0.00001), significantly reduced disease burden over that of respective controls.

**Conclusion:** We have successfully developed a novel anti-CD33 immunoPET-CT based non-invasive imaging tool for diagnosis of AML and its spatial distribution, which indicates a preferential skeletal niche.

**Translational Relevance:** Acute myeloid leukemia (AML) exhibits extensive heterogeneity in therapeutic response, even in patients sharing similar characteristics, indicating the inadequacy of currently used biopsy-based prognostic tools. There is a critical need for non-invasive quantitative imaging modality specific to AML, which is able to assess this disease in the whole body, including in extramedullary organs and to longitudinally monitor treatment response. CD33, an accepted AML biomarker is expressed on more than 85-90% of blast cells and Mylotarg (anti-CD33-ADC) is FDA approved immunotherapy for AML. In this preclinical study using anti-human CD33 mAb (^64^Cu-DOTA-anti-CD33), a CD33 PET-CT imaging modality was developed that detected AML with high sensitivity and specificity. The approach also indicated spatial heterogeneity in leukemic disease, dependent on disease burden, which warrants caution in analyzing results from single point bone marrow biopsies. Therefore, this imaging modality can be used to non-invasively detect AML in humans and also monitor treatment response, an unmet clinical need.

## Introduction

Acute myeloid leukemia (AML) is a highly aggressive hematopoietic malignancy with an extremely poor prognosis as reflected by an overall 5-year survival rate of 40%-45% in young adults and <10% in the elderly (>65 years of age) (1). Research over the past decades has helped us understand the pathobiology, classification and genomic landscape of the disease, which has resulted in improving current treatment options. Despite advances, the prognosis for elderly patients who account for the majority of new AML cases remains discouraging (2). More than 70% of elderly AML patients (> 65 years old) will die of their disease within 1 year of diagnosis and treatment (3). Therefore, new diagnostic and therapeutic approaches are necessary to improve outcomes.

Currently, the diagnostic criteria for AML is the presence of ≥20% blasts in the bone marrow or peripheral blood (4). AML diagnosis and prognosis are currently achieved by single-point bone marrow biopsies (iliac crest) followed by cytogenetics and mutation analysis. However, the iliac crest may not always be accurately representative of disease distribution within the entire body, particularly in the context of extramedullary disease, which is common in AML. Hence, there is need for new diagnostic tools that are non-invasive, specific and sensitive to AML in the whole body, including extramedullary organs, and useful to longitudinally monitor disease and treatment response.

CD33 or SIGLEC3 is a cell surface marker found on myeloid stem cells, monoblasts, myeloblasts, monocytes/macrophages, and granulocytic precursors. However, CD33 is not expressed on erythrocytes, platelets, B-cells, T-cells, or NK cells, making it a suitable myeloid marker and therefore commonly used in the diagnosis of AML. CD33 has been shown to be expressed in more than 85% of AML cells (blasts) (5), and an increased level of CD33 has been correlated with poor survival (6). Also, an anti-CD33 antibody drug conjugate (ADC) (Mylotarg) is FDA approved immunotherapy for AML. Therefore, we hypothesized that an anti-CD33 monoclonal antibody (mAb) would be an ideal candidate for immuno-positron emission tomography (PET) imaging of AML.

PET and computed tomography (CT) is an important imaging modality used in nuclear medicine. PET has an advantage in providing physiological and biochemical information to identify normal versus malignant lesions, but lacks anatomical details. However, CT provides high-resolution images with substantial anatomical details but lacks physiological information. Therefore, in the past decades, the two imaging modalities have been integrated forming PET-CT, which provides accurate diagnosis with anatomical details which is crucial in biopsy and focal radiotherapy (7).

The primary goal of this study was to evaluate an anti-CD33 mAb as a diagnostic imaging marker of AML in a preclinical mouse model. For proof of concept, we first used the murine ^64^Cu-DOTA-anti-CD33 (clone p67.6) mAb for immuno-PET-CT imaging of CD33^+^ AML cells (diagnostic) in a mouse xenograft model. However, besides detecting AML, this imaging modality interestingly also provided information about the spatial distribution of disease in the whole body. Then, with a translational objective, we produced a humanized anti-CD33 monoclonal antibody (clone Hu-M195), confimed similar CD33 PET-CT imaging of AML, and explored its therapeutic potential.

## Materials and Methods

### Antibody

Murine anti-human CD33 clone p67.6 is an IgG1 kappa monoclonal antibody (mAb) that targets human CD33-positive cells of the myeloid lineage (8,9). The hybridoma was produced in a hollow fiber bioreactor and purified by Protein G and cation exchange chromatography. Anti-human CD33 (Clone WM 53, # 555450) was obtained from BD Biosciences. Anti-human CD45 (clone: 2D1, # 368516) was obtained from BioLegend, San Diego, CA.

### Humanized anti-CD33

The murine anti-CD33 M195 mAb was humanized by CDR grafting to reduce human antimouse antibody responses (10,11). The scFv was reformatted to a human IgG1 antibody by cDNA synthesis, transiently expressed in HEK293 cells and purified by Protein A chromatography.

### Anti-human CD33 antibody DOTA and ^64^Cu conjugation

The mouse and humanized anti-human CD33 mAb were conjugated with the metal chelator 1,4,7,10-tetraazacyclododecane-N,N′,N′′,N′′′-tetraacetic acid (NHS-DOTA; Macrocyclics, Dallas, TX) as previously described (12). The detailed methods are provided in supplemental method.

### Cell lines

Human AML cells (HL60, MV4-11, Kg1a) and multiple myeloma (MM) cells MM.1S, Daudi cells were used in this study and cultured using standard tissue culture condition. The use of human samples were in accordance and approved by institutional review borad and the informed consent was obtained from patients or guardians in accordance with the Declaration of Helsinki. Detailed information about the cells and culture conditions are described in the supplementary methods section.

### Flow cytometry

Flow cytometry was used to analyze CD33 and CD45 expression in human AML (MV4-11, HL 60 and Kg1a) and multiple myeloma cells (MM1S) using mouse anti-human CD33 and CD45 antibodies. Staining and flow cytometry analysis was performed as per standard protocols. Data were acquired from a BD Fortessa cytometer and analyzed in FlowJo V 10.0 software.

### microPET-CT imaging and biodistribution studies

Mice bearing CD33^+^ AML cells (MV4-11, HL-60), CD33^-^ MM cells (MM1S) or non-leukemic control mice were injected IV with murine ^64^Cu-DOTA-anti-CD33 (100 μCi/ 10 μg), or ^64^Cu-DOTA-anti-CD33 (100 μCi/ 10 μg) + 500 μg of unlabeled anti-CD33-DOTA (1:50). For the humanized anti-CD33 mAb, human IVIg was injected (~1mg/mouse, ip) 2-3h prior to injecting humanized ^64^Cu-DOTA-anti-CD33 mAb (100 μCi/ 1μg). However, murine mAb did not necessitate human IVIg for imaging and biodistribution studies. Each group consisted of at least 5 mice; representative data for each group are presented. Static PET scans were acquired at 1 day (40 min scan), and 2 days (60 min scan, with whole body CT at 100 *μm* resolution) post-injection using InVeon PET-CT (Siemens). For biodistribution studies, mice were euthanized at 24h and/or 48h, as both time points showed activity. Various organs were obtained from control and AML leukemia-bearing mice. Wet weighs of each organ were determined, and radioactive counts from each tissue/organs were measured using a WIZARD2 automatic gamma counter (PerkinElmer). Biodistribution and imaged PET activity of ^64^Cu-DOTA-anti-CD33 was presented as the percent injected dose per gram or organ/tissue (% ID/g). MV4-11 with cold-CD33 block and CD33^-^ MM.1S served as negative controls. Post Processed PET image analysis was performed by Vivoquant (Invicro). ^64^Cu-anti-CD33-DOTA PE-CT imaging and biodistribution of MV4-11 AML bearing mice and no-leukemic control mice is representative of atleast 2 independent experiements (n=5).

### Mouse AML models: immunodeficient (NSG)

All animal experiments were carried out in accordance to the guidelines of Institutional Animal Care and Use Committee (IACUC). The NSG (NOD-*scid*IL2Rg^null^) mice were purchased from The Jackson laboratory and were in bred in city of hope animal breeding facility. The NSG mice were treated with 1-2 Gy radiation 24 hours before transplant as a preconditioning regime to ensure faster engraftment. Human AML and MM cells (1-2 x 10^6^ cells) were injected via tail vein, and engraftment was determined using bioluminescent imaging (BLI) 7-10 days post-transplant. Biodistribution and ^64^Cu-DOTA-anti-CD33 mAb imaging studies, immunotherapy interventions were performed around 2-3 weeks post-transplant.

### Murine and humanized anti-CD33 mAb blood pharmakokinetics

Humanized anti-CD33-DOTA mAb was radiolabeled as mentioned before with ^111^In (Triad, Isotope, Los Angeles, CA). The blood activity clearance of murine and humanized ^111^In-DOTA-anti-CD33 mAb was tested by collecting blood from tail veins at different points post injections (0, 2, 4, 24, 48, 72 h) and using a gamma counter. The detailed method is described in supplemental section.

### Humanized anti-CD33 and secondary (2°) Ab-ADC *in vitro* and *in vivo*

The therapeutic potential of humanized anti-CD33 mAb as an ADC was evaluated indirectly using anti-human IgG (Fc) 2° Ab conjugated to monomethyl auristatin E, (MMAE) (Moradec, LLC, San Diego, CA). The *in vitro* cytotoxicity assay was performed using a 1:1 molar ratio of anti-CD33 and 2°-ADC antibody. The primary and secondary antibody were first allowed to bind (~15 min) in complete IMDM medium and was then added to cells and incubated at 37°C, 5% CO_2_. The cytotoxicity was assayed after 5 days by measuring Annexin V and PI staining in these cells. The antibody amount used ranged from 1 ng-2.5 μg. As control, CD33^-^ Daudi cells (multiple myeloma) and secondary Ab-ADC alone (1 μg) were used.

For *in vivo* study, 2.5 μg of humanized anti-CD33 antibody and 2.5 μg of secondary Ab-ADC were mixed, incubated for 15 min at 4°C, and then resuspended in 100 μl of PBS 0.1% BSA and retro-orbitally injected into mice bearing MV4-11 leukemia on day 0 and day 3. As controls, leukemic mice were untreated or given secondary Ab-ADC alone. Disease progression was monitored using BLI at indicated time points. The whole body of the mouse was contoured, and average BLI intensity (photons/s) for each group was plotted at different time points.

### Humanized anti-CD33 radioimmunotherapy (CD33-RIT)

Humanized anti-CD33-DOTA mAb was radiolabeled as mentioned before with ^225^Ac (Oak Ridge National Laboratory, Tennesse) at a specific activity of 50 nCi/μg. For therapy, three different activities, 100 nCi, 200 nCi and 300 nCi of ^225^Ac-DOTA-anti-CD33, were used, with one single dose given to AML-bearing NSG mice (n=5). Untreated AML-bearing mice served as controls.

### Statistical analysiss

Statistical analysis was performed using ANOVA and the students-t test. The Pearsons correlation coefficients were calculated assuming a Gaussian distribution. The difference was considered significant when the p value was <0.05. All graphs and statistical analysis were generated using GraphPad Prism software V 7.2.

### Other materials and methods

Detailed methods for anti-human CD33 (clone P67.6 Hu-M195) mAb DOTA and ^64^Cu conjugation, CD33 surface expression quantification, BLI, lentiviral production and transduction in MV4-11 cells, CD33 antibody immunoreactivity, *in vitro* and *in vivo* stability, blood clearance and diagnostic accuracy calculations are provided in the Supplemental Methods section.

## Results

### Generation of ^64^Cu-DOTA anti-CD33 (p67.6) Monoclonal Antibody

For use as a immunoPET tracer, the anti-human CD33 mAb, mouse hybridoma P67.7, was produced, purified, conjugated to the metal chelate DOTA and radiolabeled with ^64^Cu. Coomassie stained SDS-PAGE gel electrophoresed under reducing conditions shows the purity, with 2 bands corresponding to the light and heavy chains (Supplemental Figure S 1A). Post-DOTA conjugation, iso-electrofocusing gel analysis shows a shift to a more acidic pH, confirming the conjugation process (Supplemental Figure S 1B). The radiolabeled ^64^Cu-DOTA-anti-CD33 mAb was analyzed by size exclusion chromatography (SEC) and the radiochromatogram showed a single peak, with a retention time corresponding to a mAb (Supplemental Figure S 1C). The immunoreactivity of DOTA-anti-CD33 mAb was evaluated by incubating the ^64^Cu-DOTA-anti-CD33 with soluble CD33-Fc antigen and analyzed by SEC. The radiochromatogram showed a faster retention time (~30 min), indicating an increase in molecular size consistent with binding to CD33 Fc soluble antigen (67-85 kDa) (Supplemental Figure S1D, E). The ^64^Cu-DOTA-anti-CD33 mAb was shown to be stable in serum both *in vitro* and *in vivo* for 48h (Supplemental Figure S1 F,G).

### CD33 cell surface expression in AML cells

The murine DOTA-anti-CD33 mAb was tested for binding and specificity by flow cytometry using CD33 positive MV4-11, HL-60 cells, Kg1a, and the CD33-negative MM.1S-GFP cells. The ^64^Cu-DOTA-anti-CD33 antibody showed immunoreactivity towards CD33-positive AML cell lines but not in CD33-negative MM.1S cells (Supplemental Figure S 2A). The immunoreactivity of the antibody was further confirmed by immunofluorescence: HL60 cells showed brighter staining than MV4-11, whereas negative control MM.1S showed no staining (Supplemental Figure S 2B). The total number of CD33 cell surface receptors per cell was determined using the BD Quantibrite PE kit (Supplemental Figure S 2C-D). The HL-60 AML cell line expressed ~55,000 cell surface CD33/cell, versus ~26,000/cell in MV4-11 Supplemental Figure S 2E).

### PET-CT imaging and biodistribution of CD33^+^ AML cells in mouse model

^64^Cu-DOTA-anti-CD33 mAb immuno PET-CT *in vivo* imaging was performed in NSG mice bearing CD33^+^ AML cells, CD33-MM cells, and non-leukemic control mice. Mice were injected with ^64^Cu-DOTA-anti-CD33 mAb (100 μCi/10 μg) 24-48h prior to PET imaging. As an additional control for specificity, the ^64^Cu-DOTA-anti-CD33 mAb (100 μCi/ 10 μg) + 500 μg of unlabeled DOTA-anti-CD33-mAb (1:50) were injected into mice bearing CD33^+^ AML cells. Bioluminescent imaging of AML and MM.1S cells in NSG mice was carried out 24 hrs prior to ^64^Cu-DOTA-anti-CD33 mAb injections. The engraftment of AML and MM cells was also measured in the femur, L-spine, and spleen using flow cytometry upon harvesting tissues/organs after imaging and biodistribution studies. PET-CT images (sagittal and coronal) clearly show CD33^+^ activity only in MV4-11 AML mice, but not in mice given cold anti-CD33, mice with MM.1S, and non-leukemic control mice (Fig. 1A, B), while the BLI images of the respective mice shows engraftment (Fig. 1C). CD33^+^ PET activity was detected in the epiphysis/metaphysis of femur, tibia, and humerus; L-spine; and pelvic bone in CD33^+^ MV4-11 bearing mice (Fig. 1D). Although BLI and flow cytometry indicated high levels of myeloma cells in MM.1S bearing mice, there was no detectable CD33^+^ activity; similarly, the mice injected with cold unlabeled CD33 antibody showed no CD33^+^ PET signal, suggesting high specificity (Fig. 1 A-C).

**Figure 1:**
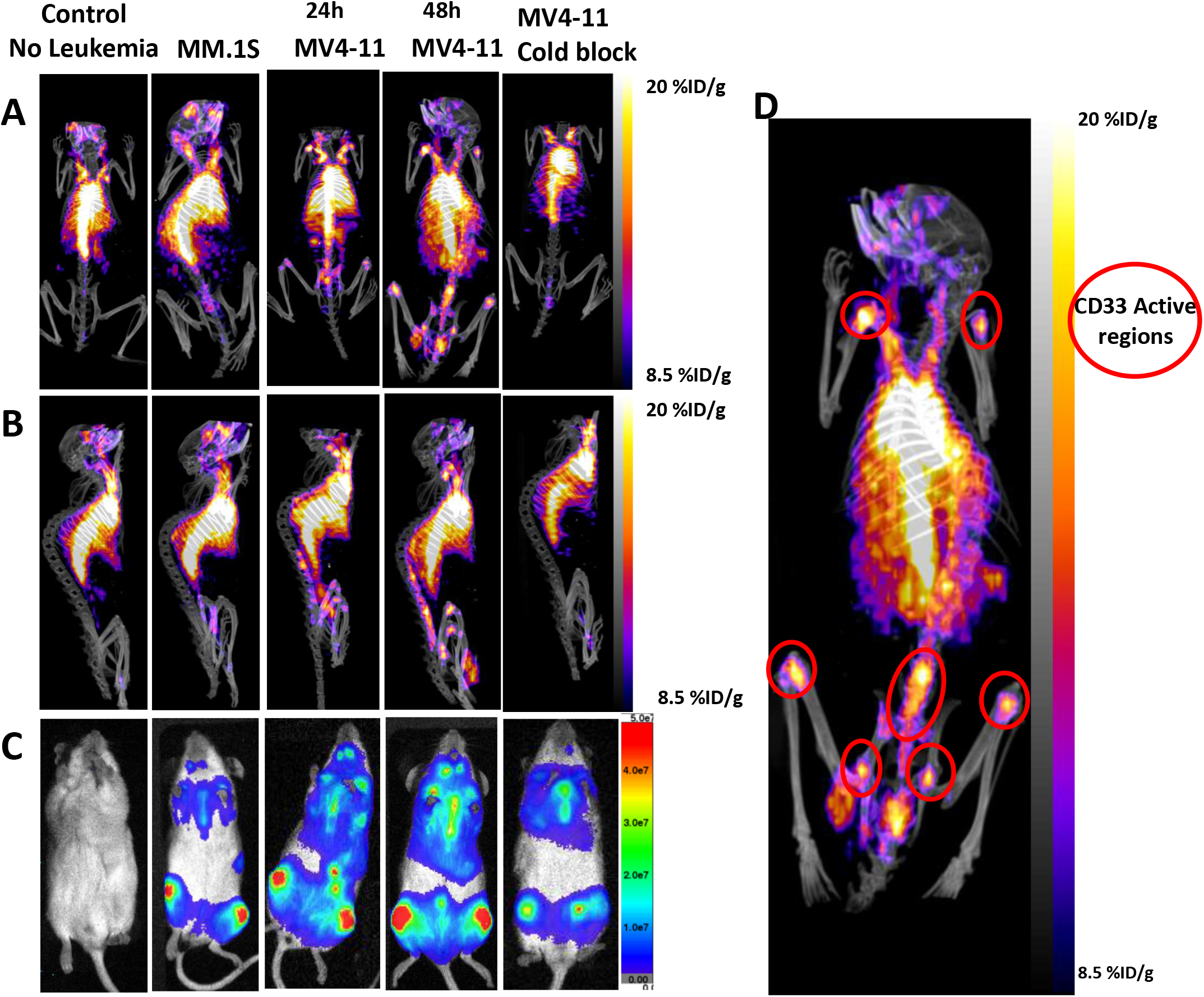
PET-CT images ^64^Cu-DOTA-anti-CD33 antibody in AML and MM bearing mice. Representative PET-CT and bioluminescence images (BLI) are shown from AML, MM bearing, and no leukemia control mice. Cu-64-anti-CD33-DOTA (100μCi/10μg) was injected into these mice via tail vein 24-48h before PET-CT imaging or biodistribution was carried out. **A**) PET-CT images (coronal) showing CD33 activity in AML bearing mice (**B**) PET-CT images (sagittal) showing CD33 activity in AML mice. (**C**) Bioluminescence (BLI) images of AML, cold blocked AML, MM.1S and no leukemia control mice. **D**) PET-CT images (coronal) of AML bearing mice highlighting CD33^+^ regions in the skeletal system. Imaged PET activity of ^64^Cu-DOTA-Anti-CD33 is presented as the percentage of the injected activity per gram of organ/tissue. Cu-64-anti-CD33-DOTA PE-CT imaging of AML bearing mice and no-leukemic control mice is representative of atleast 3 independent experiements (n=5).

Biodistribution studies were carried out in mice from respective groups by harvesting different organs/tissue 24-48h post injection of the ^64^Cu-DOTA-anti-CD33 mAb, and the activity was measured using a gamma counter. The % ID/g of tissue/organs was plotted to determine the activity. The PET-signal was similar in all groups for blood, liver, lung, heart, muscle, intestine and kidney (Fig. 2A,B). However, the % ID/g was particularly high in bones (Femur, humerus, tibia and lumber spine) in CD33^+^ bearing MV4-11 AML mice, but not in CD33^-^ MM mice, cold-blocked AML mice, or control non-leukemic mice (Fig. 2C). Therefore, MV4-11 AML mice had CD33^+^ activity in the femur, tibia, L spine and humerus. ^64^Cu-DOTA-anti-CD33 targeting was validated independently using a second AML cell line, HL-60, and similar biodistribution results were obtained (Fig. 2D, E).

**Figure 2:**
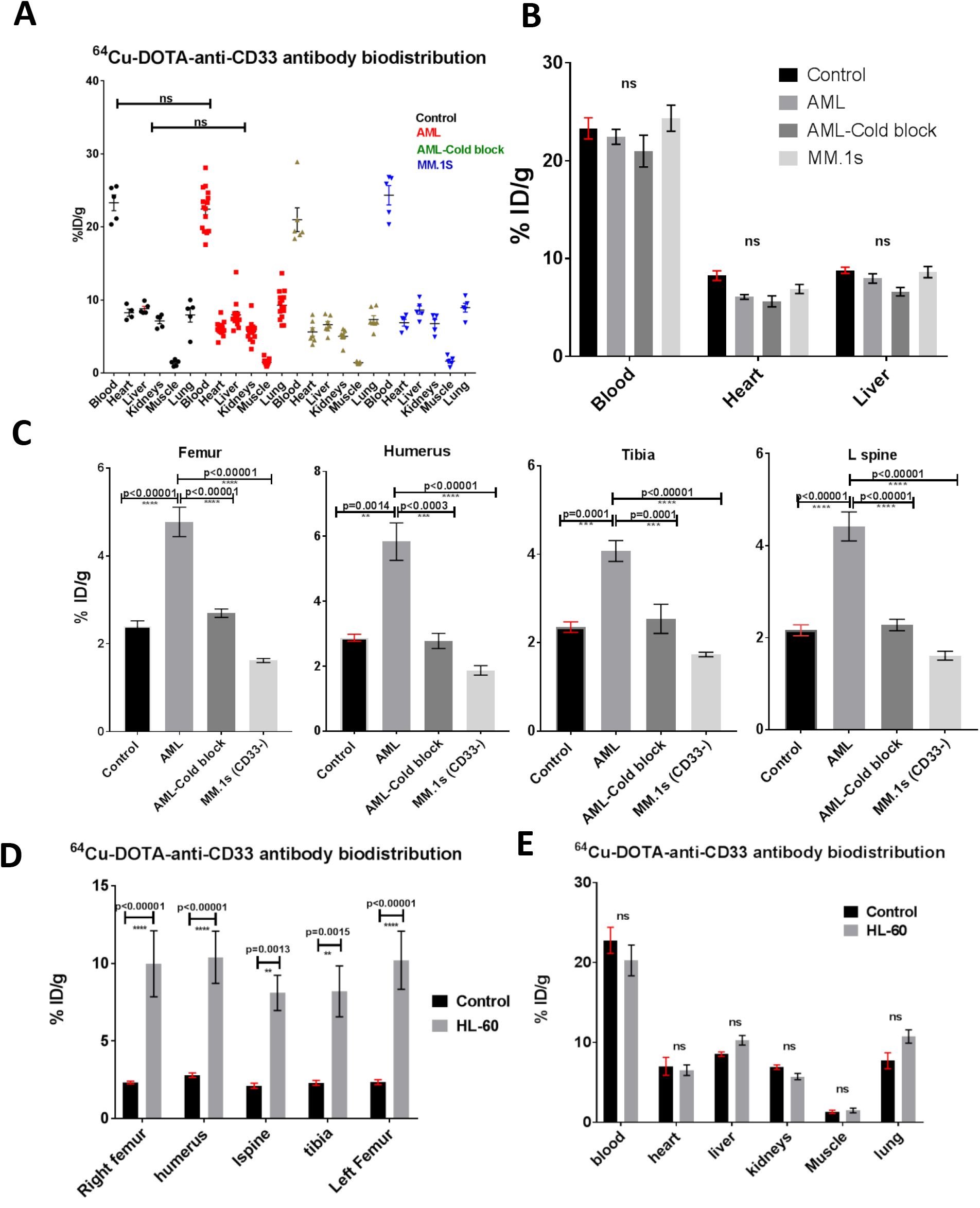
Biodistribution of ^64^Cu-DOTA-anti-CD33 antibody in AML and MM bearing mice. (**A-C**) Biodistribution of ^64^Cu-DOTA-anti-CD33 in bones and different tissues was conducted 24h/48h post injection. Plot of %ID/g of different tissues has been shown, indicating that CD33 activity is high in bones of MV4-11 mice whereas no activity was seen in CD33^-^ MM.1S, cold blocked MV4-11, or no leukemia control mice. The %ID/g between groups was insignificant for blood, heart, liver, lung and kidney. D and E) Biodistribution of ^64^Cu-DOTA-anti-CD33 in bones and soft tissues of HL-60 bearing mice in comparison to control mice. Statistical significance was determined using ANOVA/ ‘t’ test and considered significant when <0.05. Biodistribution of ^64^Cu-DOTA-Anti-CD33 was presented as the percentage of the injected activity per gram of organ/tissue.

The sensitivity and specificity were calculated from biodistribution of the ^64^Cu-DOTA-anti-CD33 mAb injected AML mice. The ROC curves showing sensitivity vs 100-specificity for biodistribution data from the femur show high sensitivity (~95%) and specificity (100%) (Fig. 3A).

**Figure 3:**
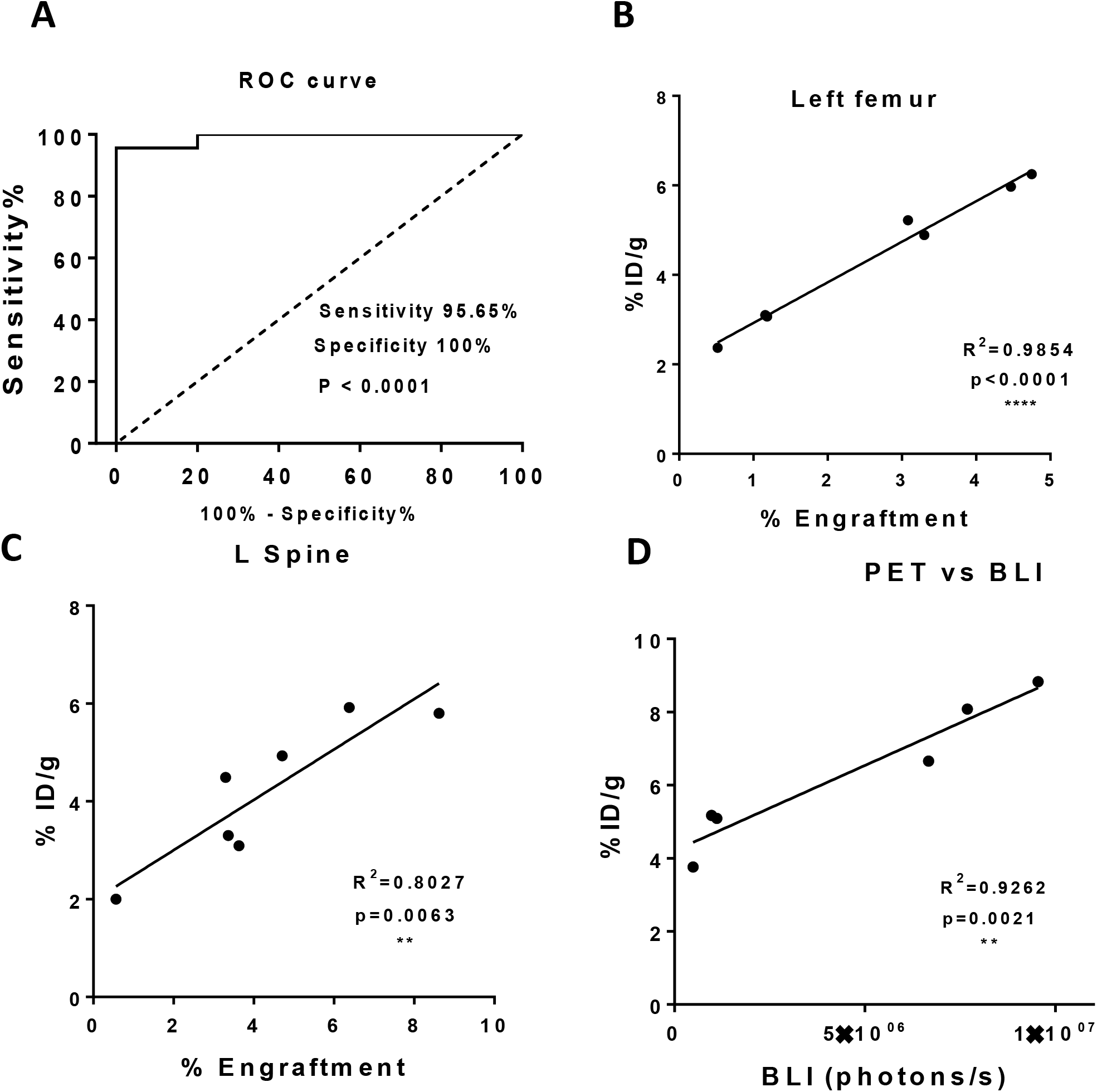
Sensitivity and specificity of anti-CD33 imaging method and correlation with leukemia engraftment. Sensitivity and specificity were calculated as mentioned in the methods. (**A**) The ROC 64 curve showing sensitivity vs 100-specificity for the Cu-DOTA-anti-CD33 imaging was generated using biodistribution data (n≥15 mice). The imaging method has a sensitivity of ~95.5% and specificity of 100 %. Therefore, this is a very reliable imaging method to detect AML. (**B and C**) A correlation curve for % engraftment vs % ID/g for femur and L-spine (n≥5 mice). A very high correlation was observed between leukemia engraftment and Cu-64-anti-CD33 activity in left femur (R^2^ = 0.9854) and L-spine (R^2^ = 0.8027). Engraftment was determined using flow cytometry. (**D**) A correlation curve for CD33 PET contour signal vs BLI signal (n≥5 mice). Strong correlation (R^2^ = 0.9262) was seen between BLI signal vs PET signal; however, the spatial resolution was high in PET whereas in BLI it was poor. The data was assumed for Gaussian distribution, and Pearson’s correlation coefficients were calculated using Prism software. The p value was calculated using two tailed t test and considered significant if <0.05.

Further, the femur and L-spine of AML bearing mice were contoured in PET-CT images. The extent of engraftment from these bones was then determined by flow cytometry of human CD45^+^ cells/ MV4-11-RFP. A high correlation was observed between PET-CT signals and percent engraftment (R^2^ value for femur and L-spine is 0.9854 and 0.8027, respectively) (Fig. 3 B, C). Additionally, the femur of AML bearing mice was contoured both in BLI and PET-CT images, and a high correlation were observed between BLI and PET-CT signals (R^2^= 0.9262) (Fig. 3D). However, spatial localization was evident in PET-CT in comparison to BLI images, which were diffuse.

### Spatial heterogeneity in AML *in vivo*

Besides detecting a CD33^+^ specific signal, anti-CD33 PET-CT imaging also indicated the spatial heterogeneity of AML. CD33^+^ PET activity was significantly heterogeneous within the femur; for example, the distal and proximal femur showed higher CD33 activity compared to the long bone area in mice with low leukemia burden; however, heightened activity in the long bone was observed when leukemic burden increased (Fig. 4A). A similar localization pattern was seen in other bones including the tibia, humerus and L-spine (Fig. 1D). CD33 activity was mostly concentrated in the proximal/distal end of the bone. Further supporting the spatial distribution of AML, in the L-spine (Fig. 4B) at areas of low leukemic burden (5-10%), the disease is strongly localized to one of the segments (L1-L5) of the L-spine. However, with increasing leukemic burden (>15%) the leukemia is distributed across the L-spine and can be seen even on the T-spine, which was initially undetectable in the low leukemia group (Fig. 4B), suggesting a preferential niche for AML during the early stages of disease progression.

**Figure 4:**
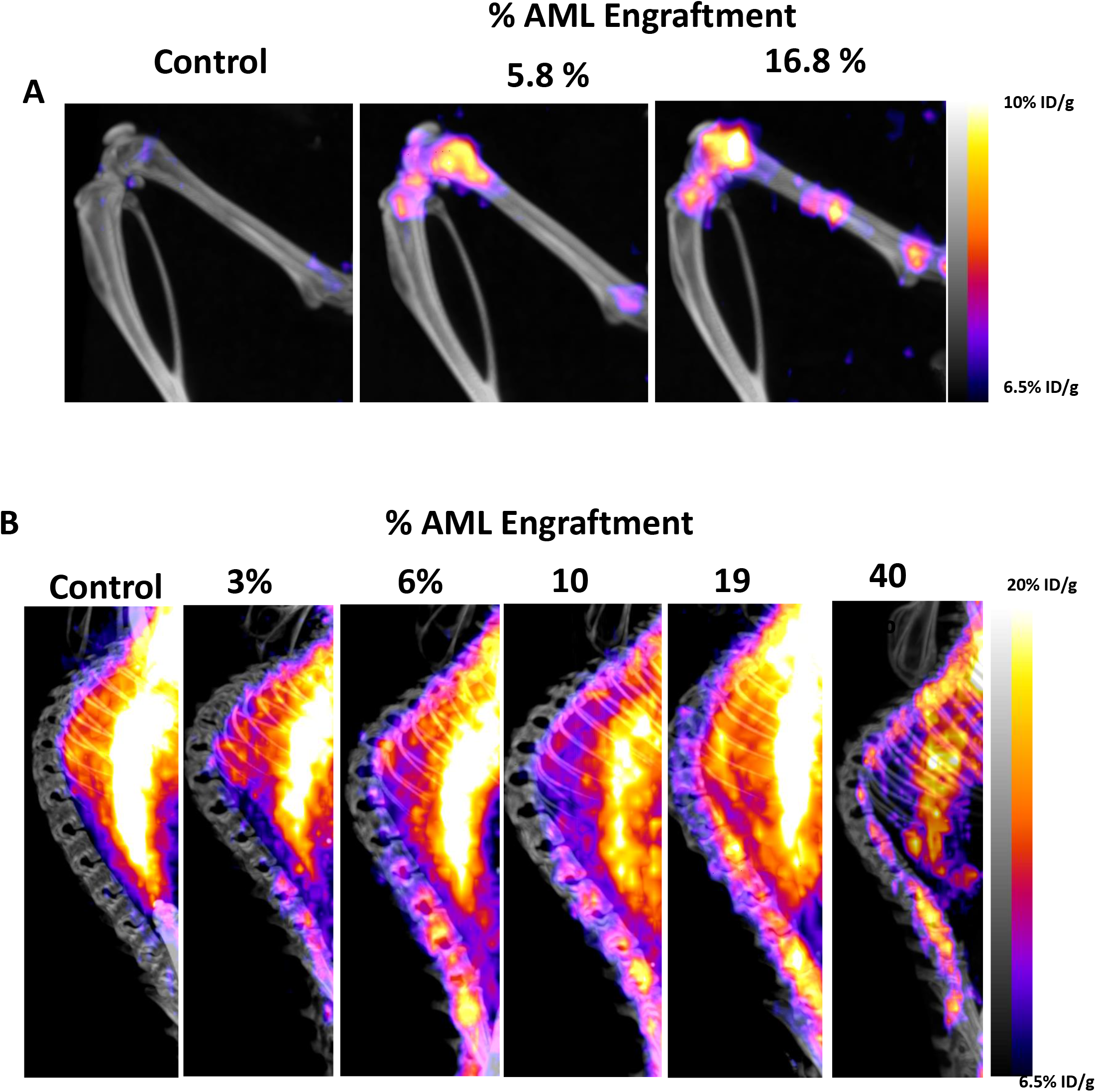
Spatial distribution of AML. (**A**) Spatial distribution of AML in femur and tibia. The PET-CT images (coronal) of representative AML bearing mice show preferential niche to the joints in the early stages of the disease. (**B**) A representative PET CT image (sagittal) of spine from AML bearing mice showing leukemia burden-dependent CD33^+^ activity. The leukemia is localized in the early disease stage (<10% engraftment); however, with increased leukemic burden (>10%), it spreads and appears systemic.

### Humanized anti-CD33 antibody detects CD33 AML

After validating the use of CD33 mAb as an imaging agent, with the intent of translating this promising technology to humans, we produced a humanized anti-CD33 mAb, as described in the methods. The radiolabeling, immunoreactivity and stability (48h, *in vivo*) were analyzed as mentioned for the murine version (Supplemental Figure S 3A-D). We tested the humanized ^64^Cu-DOTA anti-human CD33 mAb immunoreactivity and found that it bound specifically to CD33^+^ AML cell lines (MV4-11, HL-60) and human patient samples (Supplemental Figure S 4A, B). We then compared the murine and humanized CD33 mAb blood retention and the ability of the mAb to detect AML in mice bearing CD33^+^ AML cells. Significant differences between humanized and murine antibody clearance were observed at 4, 24, 48, 72 hours post injection (P=0.0021, 0.00044, 0.00073 & 0.00015 respectively) and faster blood pharmacokinetics was observed with the humanized antibody. The concentration of humanized anti-CD33 mAb in blood drops to <5% of initial concentration within 24 h post injection, while the murine antibody maintained > 25% activity of initial concentration even at 72 hours post injection (Supplemental Figure S 4C).

The humanized ^64^Cu-DOTA anti-human CD33 mAb PET-CT images (Fig. 5A,B) clearly show targeting of AML cells to the skeletal system similarly to the murine Ab, and the respective BLI image showing AML engraftment (Fig. 5C). As in the murine Ab PET imaging, humanized mAb also signaled CD33^+^ PET activity localized to the epiphysis/diaphysis of the bones, which was dependent on disease burden, indicating spatial heterogeneity (Fig. 5D). The humanized Ab also revealed CD33^+^ activity in the spleen of mouse, with ~20% AML engraftment (Fig. 5A), suggesting its ability to detect extramedullary disease. Biodistribution studies were carried out in mice from respective groups as mentioned before. There was no significant difference in PET signals in both groups in the heart, lung, muscle and kidney (Fig. 5E); however, the signal was significantly reduced in the blood while increased in liver. Also, noticably, there was reduction of the PET signal in the heart of the AML-bearing mice with the humanized antibody in comparison to the murine Ab, which may be due to lower blood activity. The % ID/g was particularly high in bones in CD33^+^ bearing AML mice, but not in control non-leukemic mice (Fig. 5F). Furthermore, the humanized mAb showed ~3-4 times higher CD33^+^ activity in bones than with murine mAb (Supplemental Figure S 4D).

**Figure 5:**
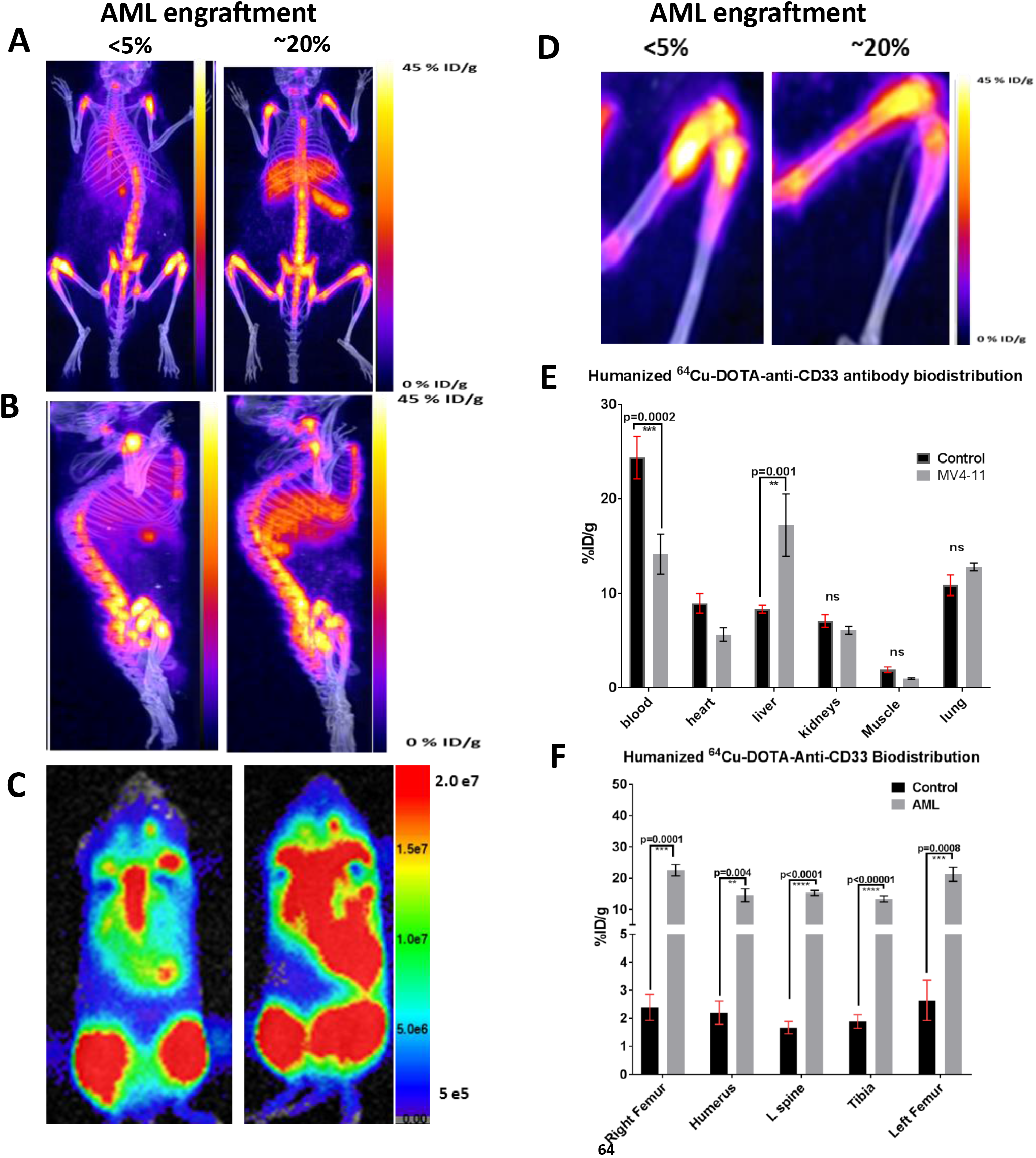
PET-CT images and Biodistribution of humanized ^64^Cu-DOTA-anti-CD33 antibody in AML bearing mice. Representative PET-CT and bioluminescence images (BLI) are shown from AML bearing, mice. Humanized ^64^Cu-anti-CD33-DOTA (100 μCi/1 μg) was injected into these mice via tail vein 24-48 h before PET-CT imaging or biodistribution was carried out. **A**) PET-CT images (coronal) showing CD33 activity in AML bearing mice (**B**) PET-CT images (sagittal) showing CD33 activity in AML mice. (**C**) Bioluminescence (BLI) images of AML bearing mice. **D)** PET-CT images (coronal: L femur) showing spatial heterogeneity in distribution of AML disease (**E, F**) Biodistribution of ^64^Cu-DOTA-anti-CD33 in bones and different tissues was conducted 24h/48h post injection. Plot of %ID/g of different tissues has been shown, indicating that CD33 activity is high in bones of MV4-11 mice whereas no activity was seen in no leukemia control mice (n=6). Statistical significance was determined using ANOVA and considered significant when <0.05. Biodistribution of ^64^Cu-DOTA-anti-CD33 was presented as the percentage of the injected activity per gram of organ/tissue.

### Humanized anti-CD33 mAb show therapeutic potential

Additionally, to asses therapeutic application of the humanized anti-CD33 mAb, as a proof of concept we have used two methods, first an ADC, an anti-human IgG (Fc) 2° Ab conjugated to MMAE, and second, ^225^Ac-DOTA anti-human CD33 radioimmunotherapy (CD33-RIT). The *in vitro* cytotocxicity assay on MV4-11 cells using anti-CD33 mAb + 2°-ADC Ab (CD33-2°-ADC) was carried out as mentioned in methods. As control, 2°-ADC Ab alone (1 μg) was used, which showed no significant cell killing over untreated cells, while the same concentration of CD33-2°-ADC yielded ~55% cell killing (Fig. 6A), and maximum (>95%) cell killing was observed at 55 nM (2.5 μg). Multiple myeloma cells (Daudi) were used to show that the CD33-2°-ADC had no significant cell killing on the CD33^-^ cell line, indicating specificity. For *in vivo* study, we used 2.5 μg CD33-2°-ADC, which showed maximum cell killing in the *in vitro* study. The MV4-11 AML bearing mice treated with CD33-2°-ADC showed decreased BLI intensity 24 days post treatement than in mice treated with 2°-ADC alone (p=0.02) or untreated AML-bearing mice (p=0.0008) (Fig. 6B, Supplemental Figure S 5A).

**Figure 6:**
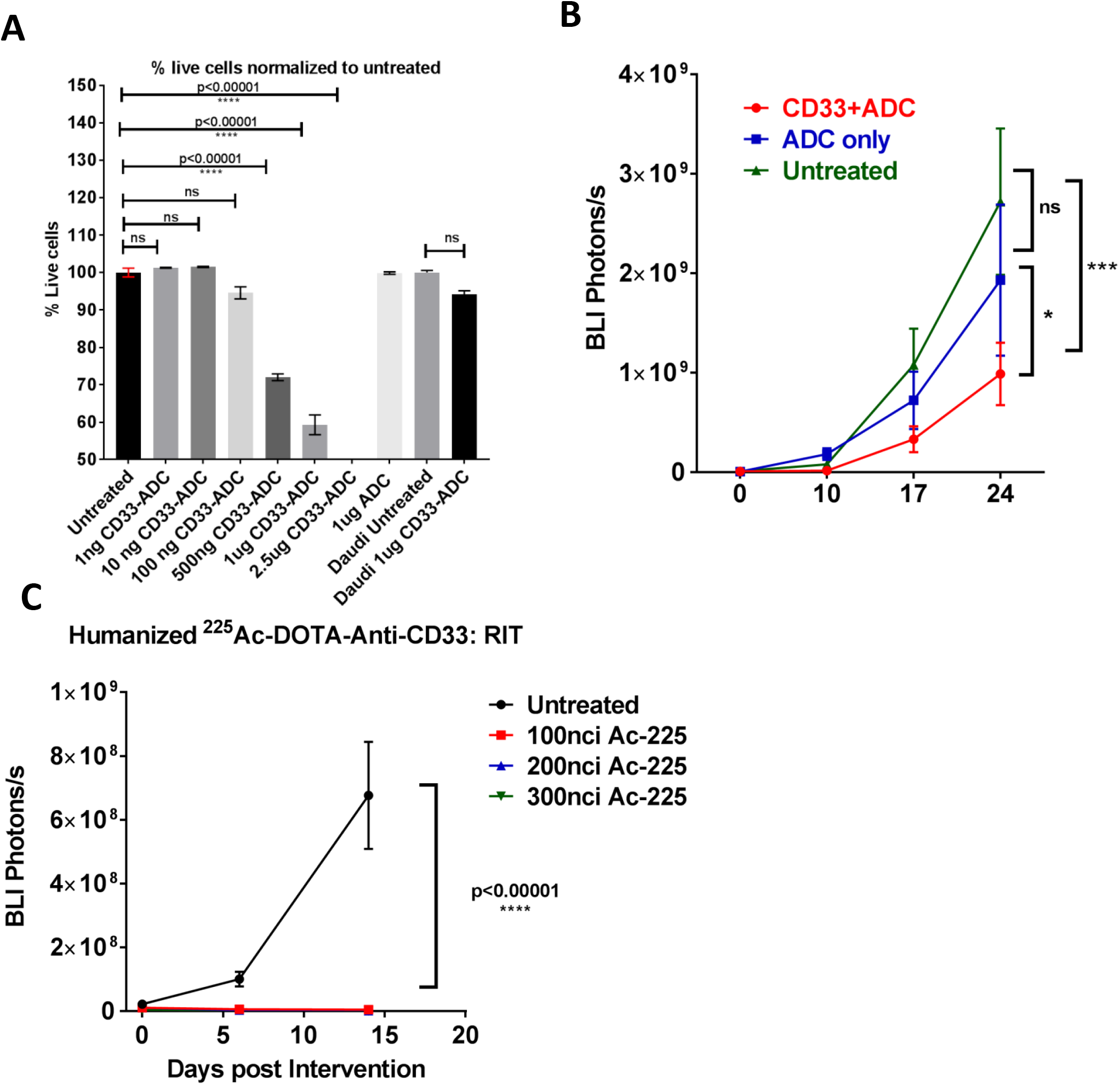
Humanized anti-CD33 mAb antibody therapeutic potential. The humanized anti-CD33 mAb therapeutic potential *in vitro* and *in vivo* as an ADC and RIT was explored. A) *In vitro* cell killing of MV4-11 cells by anti-CD33 + ADC. Significant cell killing over untreated cells was seen from 500 ng −2.5 μg of Ab. **B)** AML bearing mice was treated with Anti-CD33 mAb + 2° Ab (anti-Human Fc) conjugated to an MMAE. Untreated mice or 2° Ab-ADC only treated mice was used as controls. The BLI intensity (photons/s) of whole mouse was calculated and plotted at day 24 days post intervention. The mice treated with the anti-CD33 + 2° Ab-ADC combination reduced disease burden significantly in comparison to ADC only treated mice (p=0.02) or untreated mice (p=0.0008) (n≥4). **C)** Three different activity of humanized ^225^Ac-DOTA-anti-CD33 (100nci, 200nci, 300nci) was injected into mice bearing AML. All activity significantly reduced leukemia burden (p<0.00001) as measured by BLI intensity (photons/s) at 14 days post intervention in comparison to untreated AML bearing mice (n=5). Statistical significance was determined using ANOVA and considered significant when <0.05.

CD33-RIT was carried out using three different ^225^Ac activities (100 nCi, 200 nCi and 300 nCi). Similar to the ADC study, CD33-RIT treated mice showed a significant decrease (p<0.00001) in leukemia burden 14 days post intervention (Fig. 6C, Supplemental Figure S 5B). These results suggest that the humanized anti-CD33 antibody can be used for therapy.

Currenlty, a cGMP grade humanized antibody is under preparation for a planned phase I/II clinical trial of PET-CT imaging of AML. Efforts are also in progress for generating a primary anti-CD33 mAb-ADC to explore anti-CD33 immunotherapy using this humanized antibody and to monitor treatement response using anti-CD33 PET-CT imaging.

## Discussion

We have developed a non-invasive anti-CD33 immunoPET-CT imaging method for the *in vivo* detection of AML disease with high sensitivity and specificity. Diagnostic and prognostic markers facilitate stratifying patients for treatment management. However, the current clinically approved diagnostic method is invasive, relying on single point (iliac crest) biopsies, which may 1) not always be representative of the actual disease state, and 2) limits the number of times it may be performed. Therefore, in leukemia non-invasive PET-imaging using PET tracers including F18-FDG (metabolic activity) and F18-FLT (cell proliferation) have been tested for diagnosis and monitoring treatment response (13-15,16). Although FDG-PET has shown some success in diagnosing extramedullary disease in AML(17), it has also yielded highly inconsistent results because of changes in the metabolic activity of normal/tumor cells post treatment (18,19). Also, a recent study indicated that FLT-PET could identify the risk of early relapse in the spine prior to evidence of relapse by detection of minimal residual disease (MRD) (20). However, these PET tracers are non-specific, they detect metabolically active or highly proliferating cells, which could be effected post treatment, unlike CD33-PET imaging which specifically detects cells expressing CD33, an accepted biomarker for AML. Studies have shown that even relapse AML blast post anti-CD33 therapy have CD33 expression (21,22). Therefore, CD33-based imaging is expected to improve detection of AML with high specificity in the whole body. This approach may also provide a blueprint for selecting biopsy sites (image-guided) that would be greatly useful for early detection and also in longitudinal monitoring of treatment response.

In this study, the ^64^Cu-DOTA-anti-CD33 mAb PET was used to detect AML *in vivo,* showing very high specificity and sensitivity to CD33^+^ AML. The spatial information was achieved with whole body CT based 3D anatomical imaging. Notably, a spatial heterogeneity in the distribution of AML within the skeletal regions was observed, which would not be established with point biopsies. For example, CD33 activity was especially prominent in the L-spine and epiphysis/diaphysis of femurs, indicating a preferential skeletal niche for the disease at early stages. However, when the disease progresses, this spatial heterogeneity was reduced and the disease appeared to be more systemic. This finding suggests, contrary to previous beliefs that leukemia was a systemic disease, it appears that leukemia may be initiated in multiple preferential niches, characterized by multifocal disease before becoming more systemic. This imaging observation is warranting future investigation to understand the role of micro-enivironment favoring preferential disease localization. However, one small clinical study suggested that a heterogeneous distribution of leukemia may indicate resistance to chemotherapy, as shown through FLT-PET-based imaging of cell proliferation (23). Therefore, future clinical studies may be designed to assess the prognostic value of this anti-CD33 oncoPET imaging. As immuno-PET combines the sensitivity and resolution of a PET imaging system with the high specificity of a monoclonal antibody, it may represent what is described as a “comprehensive immunohistochemical staining *in vivo*” (24), therefore guiding precise locations for biopsies. Also, precisely specifying sites of disease could yield valuable information about the tumor in terms of location, phenotype, treatment susceptibility and response (25). The spatial heterogeneity of leukemia also suggests caution should be taken when interpreting data from single-point biopsies.

Several investigational CD33-targeted therapeutics have been developed for AML (26), including the recently FDA-approved anti-CD33 monoclonal antibody-drug conjugate gemtuzumab ozogamicin (GO, Mylotarg) (27), bispecific antibodies such as CD33/CD3 antibodies (28), CD33/CD123-directed CAR T cells (29). Although positive results have been reported from clinical trials of CD33-targeting drugs, dose limiting toxicities such as hepatotoxicities (30) during treatment are of concern. Therefore, an imaging tool as presented here would help determine optimal doses as well as monitoring treatment response non invasively, providing an opportunity to improve treatment efficacy while decreasing toxicities.

As the initial study was conducted using a murine anti-CD33 mAb, clone p67.6, for translational purposes, a humanized anti-CD33 monoclonal antibody (Hu-M195) was generated. We carried out preclinical PET-CT imaging using humanized anti-CD33 mAb and showed that targeting was significantly improved over that of the murine version. Currently, cGMP grade mAb has been made for future phase I/II clinical trials. The Hu-M195 mAb labelled with ^131^I has been shown to be useful for whole-body-gamma imaging of patients with myeloid leukemia (31). However, this modality is planar or 2D imaging and is hence qualitative and lacks 3D quantitative information with spatial heterogeneity assessment of the disease. In contrast, ^64^Cu-anti-CD33 PET-CT involves 3D imaging, which is quantitative and provides spatial information, resulting in a signifcanlty improved diagnostic imaging.

We further show that targeting CD33 using CD33-2°-ADC or CD33-RIT resulted in reduced disease burden, emphasizing its therapeutic potential. Such approaches are very effective against receptors like CD33, which is endocytosed after Ab binding, enabling the cytotoxic payload to internalize and result in cell killing (32,33). Futhermore, after CD33 antibody therapy, blasts in relapsed/refractory disease continue to express CD33, suggesting that AML cells may not use downregulation of CD33 as an epigenetic “escape mechanism” (22), in contrast to the use of anti-CD38 mAb daratumumab which may result in relapse due to a CD38^-^ myeloma clone (34). Therefore, CD33 PET imaging would be useful in monitoring treatment response post intervention, including CD33 targeted treatment. We would futher validate this approach preclinically using the primary humanized anti-CD33-ADC (which is in development) or RIT post CD33-directed therapy.

There are some limitations of this study. Although CD33 is expressed in all patients on at least a subset of AML blasts, high inter-patient variability in terms of level of expression (> 2 fold) is known (6,35,36). This differential expression is not random but correlated with the presence of particular mutations; for example, very high CD33 expression has been shown in AML with NPM1 and FLT3/ITD mutations, whereas lower expression has been associated with core binding factor translocations (6,36,37). A properly controlled imaging study with subset (mutation) analysis will be essential to understand whether specific mutations impact imaging intensity and/or heterogeneity.

In conclusion, to the best of our knowledge, this is the first preclinical report of an anti-CD33-PET-CT imaging modality to successfully detect AML *in vivo*. The imaging tool may be used for diagnosis as well as monitoring treatment response in the whole body, including extramedullary organs. The molecular imaging method also detected heterogeneity in the spatial distribution of the AML, warranting caution in interpreting results from single-point biopsies. Future studies to understand the relationship between genomic mutation (like NPM1, FLT3-ITD etc.) and spatial heterogeneity (as observed in the CD33 PET-CT imaging) would be beneficial in the diagnosis and prognosis of AML. In summary, anti-CD33 PET-CT is a viable AML imaging method with significant translational potential.

## Acknowledgments

We appreciate administrative support at our institutions. Research reported in this publication included work performed by the Small Animal Imaging Core for PET-CT imaging and imaging precision radiation delivery system supported by the National Cancer Institute of the National Institutes of Health under award number P30CA033572 and partly supported by National Institutes of Health grant 1R01CA154491-01. The content is solely the responsibility of the authors and does not necessarily represent the official views of the National Institutes of Health.

## Authorship

Contribution: SSM, DZ, JB, BK, LEP, MO, PV, IN, JC, KP, NB, AM, TE and JM performed experiments. SSM, JB, DZ and SKH analyzed the data and made the figures. SKH, SSM, JB, DZ, AS, JYW, JR, DAV, DC, C-CC, JES, PJY and JS were involved in designing the research. SSM and SKH wrote the paper. JS and PJY helped in editing the manuscript. All authors reviewed the paper.

## Abbreviations

3D: three dimensional
^64^Cu: copper64
AML: acute myeloid leukemia
BM: bone marrow
CBCT: cone beam CT
CT: computed tomography
FDG: fluorodeoxyglucose
FLT: flourothymidine
FLT3/ITD: FLT3 internal tandem duplication
fTMI: functional total marrow irradiation
Gy: Gray
HSCT: hematopoietic stem cell transplant
MM: multiple myeloma
NHS-DOTA: 1,4,7,10-tetraazacyclododecane-N,N’,N’’,N’’’-tetraacetic acid
NPM1: nucleophosmin 1
NSG: NOD-*scid*IL2Rg^null^
PET: positron emission tomography
SIGLEC3: Sialic Acid-Binding Ig-Like Lectin 3
TBI: total body irradiation
ADC: Antibody drug Conjugate
RIT: Radioimmunotherapy
MMAE: monomethyl auristatin E.

## References

1. Hassan C, Afshinnekoo E, Li S, Wu S, Mason CE. Genetic and epigenetic heterogeneity and the impact on cancer relapse. Experimental hematology 2017;54:26–30 doi 10.1016/j.exphem.2017.07.002.

2. Shah A, Andersson TM, Rachet B, Bjorkholm M, Lambert PC. Survival and cure of acute myeloid leukaemia in England, 1971-2006: a population-based study. British journal of haematology 2013;162(4):509–16 doi 10.1111/bjh.12425.

3. Mayer AT, Natarajan A, Gordon SR, Maute RL, McCracken MN, Ring AM, et al. Practical Immuno-PET Radiotracer Design Considerations for Human Immune Checkpoint Imaging. Journal of Nuclear Medicine 2017;58(4):538–46 doi 10.2967/jnumed.116.177659.

4. Dohner H, Estey EH, Amadori S, Appelbaum FR, Buchner T, Burnett AK, et al. Diagnosis and management of acute myeloid leukemia in adults: recommendations from an international expert panel, on behalf of the European LeukemiaNet. Blood 2010;115(3):453–74 doi 10.1182/blood-2009-07-235358.

5. Ehninger A, Kramer M, Röllig C, Thiede C, Bornhäuser M, von Bonin M, et al. Distribution and levels of cell surface expression of CD33 and CD123 in acute myeloid leukemia. Blood Cancer Journal 2014;4(6):e218 doi 10.1038/bcj.2014.39.

6. Pollard JA, Alonzo TA, Loken M, Gerbing RB, Ho PA, Bernstein ID, et al. Correlation of CD33 expression level with disease characteristics and response to gemtuzumab ozogamicin containing chemotherapy in childhood AML. Blood 2012;119(16):3705–11 doi 10.1182/blood-2011-12-398370.

7. Griffeth LK. Use of PET-CT scanning in cancer patients: technical and practical considerations. Proceedings (Baylor University Medical Center) 2005;18(4):321–30.

8. Appelbaum FR, Matthews DC, Eary JF, Badger CC, Kellogg M, Press OW, et al. The use of radiolabeled anti-CD33 antibody to augment marrow irradiation prior to marrow transplantation for acute myelogenous leukemia. Transplantation 1992;54(5):829–33.

9. van der Jagt RH, Badger CC, Appelbaum FR, Press OW, Matthews DC, Eary JF, et al. Localization of radiolabeled antimyeloid antibodies in a human acute leukemia xenograft tumor model. Cancer research 1992;52(1):89–94.

10. Caron PC, Co MS, Bull MK, Avdalovic NM, Queen C, Scheinberg DA. Biological and immunological features of humanized M195 (anti-CD33) monoclonal antibodies. Cancer research 1992;52(24):6761–7.

11. Co MS, Avdalovic NM, Caron PC, Avdalovic MV, Scheinberg DA, Queen C. Chimeric and humanized antibodies with specificity for the CD33 antigen. Journal of immunology (Baltimore, Md: 1950) 1992;148(4):1149–54.

12. Caserta E, Chea J, Minnix M, Viola D, Vonderfecht S, Yazaki P, et al. Copper 64-labeled daratumumab as a PET-CT imaging tracer for multiple myeloma. Blood 2018;131(7):741–5 doi 10.1182/blood-2017-09-807263.

13. Stölzel F, Röllig C, Radke J, Mohr B, Platzbecker U, Bornhäuser M, et al. (18)F-FDG-PET-CT for detection of extramedullary acute myeloid leukemia. Haematologica 2011;96(10):1552–6 doi 10.3324/haematol.2011.045047.

14. Arimoto MK, Nakamoto Y, Nakatani K, Ishimori T, Yamashita K, Takaori-Kondo A, et al. Increased bone marrow uptake of 18F-FDG in leukemia patients: preliminary findings. SpringerPlus 2015,’4:521 doi 10.1186/s40064-015-1339-2.

15. Han EJ, Lee BH, Kim JA, Park YH, Choi WH. Early assessment of response to induction therapy in acute myeloid leukemia using (18)F-FLT PET/CT. EJNMMI research 2017;7(1):75 doi 10.1186/s13550-017-0326-8.

16. Magome T, Froelich J, Holtan SG, Takahashi Y, Verneris MR, Brown K, et al. Whole-Body Distribution of Leukemia and Functional Total Marrow Irradiation Based on FLT-PET and Dual-Energy CT. Molecular Imaging 2017;16:1536012117732203 doi 10.1177/1536012117732203.

17. Cribe AS, Steenhof M, Marcher CW, Petersen H, Frederiksen H, Friis LS. Extramedullary disease in patients with acute myeloid leukemia assessed by 18F-FDG PET. European journal of haematology 2013;90(4):273–8 doi 10.1111/ejh.12085.

18. Riad R, Omar W, Sidhom I, Zamzam M, Zaky I, Hafez M, et al. False-positive F-18 FDG uptake in PET-CT studies in pediatric patients with abdominal Burkitt’s lymphoma. Nuclear medicine communications 2010;31(3):232–8 doi 10.1097/MNM.0b013e328334fc14.

19. Long NM, Smith CS. Causes and imaging features of false positives and false negatives on F-PET-CT in oncologic imaging. Insights into imaging 2011;2(6):679–98 doi 10.1007/s13244-010-0062-3.

20. Williams KM, Holter-Chakrabarty JL, Lindenberg L, Adler S, Chai A, Kurdziel K, et al. Novel PET Imaging with Fluorothymidine (FLT) Predicts Relapse Quantitatively at Day 28 Post Transplantation in Patients with Acute Leukemia. Biology of Blood and Marrow Transplantation 2016;22(3):S213–S4 doi 10.1016/j.bbmt.2015.11.611.

21. Linenberger ML. CD33-directed therapy with gemtuzumab ozogamicin in acute myeloid leukemia: progress in understanding cytotoxicity and potential mechanisms of drug resistance. Leukemia 2005;19(2):176–82 doi 10.1038/sj.leu.2403598.

22. Chevallier P, Robillard N, Ayari S, Guillaume T, Delaunay J, Mechinaud F, et al. Persistence of CD33 expression at relapse in CD33(+) acute myeloid leukaemia patients after receiving Gemtuzumab in the course of the disease. British journal of haematology 2008; 143(5):744–6 doi 10.1111/j.1365-2141.2008.07153.x.

23. Vanderhoek M, Juckett MB, Perlman SB, Nickles RJ, Jeraj R. Early assessment of treatment response in patients with AML using [(18)F]FLT PET imaging. Leukemia research 2011;35(3):310–6 doi 10.1016/j.leukres.2010.06.010.

24. Verel I, Visser GWM, van Dongen GA. The Promise of Immuno-PET in Radioimmunotherapy. Journal of Nuclear Medicine 2005; 46(1 suppl):164S–71S.

25. Knowles SM, Wu AM. Advances in Immuno-Positron Emission Tomography: Antibodies for Molecular Imaging in Oncology. Journal of Clinical Oncology 2012;30(31):3884–92 doi 10.1200/jco.2012.42.4887.

26. Walter RB. Investigational CD33-targeted therapeutics for acute myeloid leukemia. Expert opinion on investigational drugs 2018;27(4):339–48 doi 10.1080/13543784.2018.1452911.

27. Castaigne S, Pautas C, Terre C, Raffoux E, Bordessoule D, Bastie JN, et al. Effect of gemtuzumab ozogamicin on survival of adult patients with de-novo acute myeloid leukaemia (ALFA-0701): a randomised, open-label, phase 3 study. Lancet (London, England) 2012;379(9825):1508–16 doi 10.1016/s0140-6736(12)60485-1.

28. Gasiorowski RE, Clark GJ, Bradstock K, Hart DN. Antibody therapy for acute myeloid leukaemia. British journal of haematology 2014;164(4):481–95 doi 10.1111/bjh.12691.

29. Petrov JC, Wada M, Pinz KG, Yan LE, Chen KH, Shuai X, et al. Compound CAR T-cells as a double-pronged approach for treating acute myeloid leukemia. Leukemia 2018;32(6):1317–26 doi 10.1038/s41375-018-0075-3.

30. Godwin CD, McDonald GB, Walter RB. Sinusoidal obstruction syndrome following CD33-targeted therapy in acute myeloid leukemia. Blood 2017;129(16):2330–2 doi 10.1182/blood-2017-01-762419.

31. Caron PC, Jurcic JG, Scott AM, Finn RD, Divgi CR, Graham MC, et al. A phase 1B trial of humanized monoclonal antibody M195 (anti-CD33) in myeloid leukemia: specific targeting without immunogenicity. Blood 1994;83(7):1760–8.

32. Walter RB, Raden BW, Zeng R, Hausermann P, Bernstein ID, Cooper JA. ITIM-dependent endocytosis of CD33-related Siglecs: role of intracellular domain, tyrosine phosphorylation, and the tyrosine phosphatases, Shp1 and Shp2. Journal of leukocyte biology 2008;83(1):200–11 doi 10.1189/jlb.0607388.

33. Giles F, Estey E, O’Brien S. Gemtuzumab ozogamicin in the treatment of acute myeloid leukemia. Cancer 2003;98(10):2095–104 doi 10.1002/cncr.11791.

34. Minarik J, Novak M, Flodr P, Balcarkova J, Mlynarcikova M, Krhovska P, et al. CD38-negative relapse in multiple myeloma after daratumumab-based chemotherapy. European journal of haematology 2017;99(2):186–9 doi 10.1111/ejh.12902.

35. Hauswirth AW, Florian S, Printz D, Sotlar K, Krauth M-T, Fritsch G, et al. Expression of the target receptor CD33 in CD34+/CD38-/CD123+ AML stem cells. European Journal of Clinical Investigation 2007;37(1):73–82 doi doi:10.1111/j.1365-2362.2007.01746.x.

36. Krupka C, Kufer P, Kischel R, Zugmaier G, Bögeholz J, Köhnke T, et al. CD33 target validation and sustained depletion of AML blasts in long-term cultures by the bispecific T-cell-engaging antibody AMG 330. Blood 2013 doi 10.1182/blood-2013-08-523548.

37. De Propris MS, Raponi S, Diverio D, Milani ML, Meloni G, Falini B, et al. High CD33 expression levels in acute myeloid leukemia cells carrying the nucleophosmin (<em>NPM1</em>) mutation. Haematologica 2011;96(10):1548–51 doi 10.3324/haematol.2011.043786.

