## Supplemental Figures for "AML ImmunoPET: ^64^Cu-DOTA-anti-CD33 PET-CT imaging of AML in Xenograft mouse model"

### Supplemental methods:

#### AML and MM Cells

Human AML (MV4-11 cells, HL-60 cells, Kg1a) were purchased from the *American* Tissue Culture Collection (U.S.A.). GFP-Luciferase positive (Gfp/Luc) MM.1S MM cells were provided by Dr. Xiuli Wang (City of Hope, CA). Human AML cell lines were cultured in IMDM medium; all media were supplemented with 10% fetal bovine serum. Cells were cultured at 5% $CO_2$, 37°C in a humidified incubator. MV4-11 and HL-60 cells were 100% positive for CD33 while Kg1a had <30% cells positive for CD33. Therefore for further studies we only used MV4-11 and HL-60 AML cells (Fig. 2A). Mycoplasma testing by PCR was performed every three months on cell lines in culture. The cell lines were used within 10-12 passages for *in vivo* studies.

#### Anti-CD33-DOTA immunoreactivity

All radiolabeled antibodies were analyzed for immunoreactivity to soluble CD33 by a liquid phase assay incubating the radiolabeled protein with 20 equivalents by the mass of purified CD33 at 37°C for 15 min. The resultant solution was analyzed by HPLC-SEC using a Superose 6 10/300 GL column (GE Healthcare). Anti-CD33 immunoreactivity was determined by integrating the area on the HPLC radiochromatogram and calculating the percentage of radioactivity shifting to higher molecular weights, consistent with binding to CD33 Fc antibody (67-85 kDa).

#### Anti-human CD33 antibody DOTA and $^{64}$Cu conjugation

Briefly, 5 mg of antibody was buffer exchanged into sodium bicarbonate buffer, conjugated to NHS-DOTA, (DOTA:mAb molar ratio of 30:1, 1 hour at room temperature), buffer exchanged into 0.25 M ammonium acetate pH 7.0, and concentrated to >5 mg/ml.

Anti-CD33-DOTA mAb was radiolabeled with $^{64}$Cu (Mallinckrodt Institute of Radiology, Washington University School of Medicine) at a specific activity of 10 µCi/mg in 0.25 M $NH_4OAc$, pH 5.0 for 45 min at 43˚C, chased with 1 mM diethylenetriamine pentaacetic acid (DTPA), following which the $^{64}$Cu-DOTA-anti-CD33 conjugate was purified on a size-exclusion Superdex-200 preparative column (GE Healthcare Life Sciences).

The anti-human CD33 mAb, DOTA-anti-human CD33 mAb and DOTA-anti-human CD33 mAb vialed product were analyzed by SDS-PAGE (4-12% gradient polyacrylamide) and isoelectric focusing, and the gels were stained with Coomassie Brilliant Blue for visualization.

#### Stability Studies

Protein stability studies were performed on $^{64}$Cu DOTA-anti-CD33 incubated in fresh mouse serum at 37°C. Aliquots were analyzed on an HPLC SEC Superose 6 column at 4, 24 and 48h.

#### Dylight-488 conjugation of Clone P67.6

The P67.6 antibody was conjugated to Dylight-488 as per manufacturer's protocol (Abcam Dylight® 488 Fast Conjugation Kit (ab201799). Briefly, to 100µg of antibody (in 100µl) 10µl of modifier reagent (1µl/10µl antibody mix, [1:10 v/v]) was added. The mixture was then added to Dylight-488 conjugation reagent and incubated for 15 min in the dark. The reaction as stopped by adding 10 µL (1µl/10 µl antibody mix, (1:10 v/v)) of quenching reagent.

#### CD33 expression

The cell surface expression of CD33 on AML cells and multiple myeloma cells was determined using BD QuantiBRITE PE system as per the manufacturer's instruction. Briefly, 1 x $10^6$ cells were stained with PE-conjugated anti-CD33 MoAb (BD Biosciences) and 20,000 events were acquired for each sample using BD Fortessa cytometer, and data were analyzed using FlowJo V10.0. The BD QuantiBRITE PE contains four sets of beads with PE-molecules covalently attached to beads at four different levels. Geometric means (MFI) of all four beads were determined and using lot-specific values for the PE molecules per bead (provided in each BD Quantibrite PE kit box) $\log_{10}$ for geometric mean (MFI) and for the PE-molecules per bead was calculated. Then, a linear regression was plotted for $\log_{10}$ PE molecules per beads against $\mathrm{Log}_{10}$ geometric mean using y=mx+c equation, where *y* equals $\mathrm{Log}_{10}$ geometric mean and *x* equals Log10 PE molecules per bead. The CD33 molecules per cell on AML cells were determined by substituting the Log geometric means (y) in the equation and solved for Log PE molecules per cell (x). Anti-log of x resulted in the total number of CD33 molecules per cell.

#### Immunofluorescence staining of CD33 on AML cells

The AML and MM cells were stained with DOTA-anti-CD33-Dylight-488 mAb and immunofluorescence imaging was carried out using standard procedure and images were acquired using Zeiss AxioObserver Z1 florescent microscope. Briefly, ~ 100,000 Cells were stained with anti-CD33 mAb for 30 minutes and the stained cells were cytospun onto glass slide using Cytospin™4 (Thermo Fisher Scientific) for 5 min at 700 rpm. The samples were fixed in 4% PFA at room temperature for 5 min, cover slipped with ProLong™ Gold antifade mountant with DAPI (Thermo) and imaged. Unstained cells were used to set background, and all images were obtained at 20X magnification.

### Lentivirus preparation and transduction into MV4-11 cell line

MI-Luciferase-IRES-mCherry was a gift from Xiaoping Sun (Addgene plasmid # 75020) [1]. MI-Luciferase-IRES-mCherry plasmid (10µg) co-expressing mCherry and luciferase along with VSVG envelope and CMV packaging vectors were transfected into HEK293T cells, and supernatant was collected at 72h. The supernatant containing viral particles was mixed with 5X PEG (SBI system Biosystems, Palo Alto, CA) and kept overnight at 4°C in the rotor. The next day, the supernatant was centrifuged at 2000 rpm for 10 mins to collect viral particles in a pellet, which was re-suspended into serum free X-VIVO™ 10 media (Lonza) and stored at -80°C until further use. The titer of the viral particles was quantified using transducing HEK cells. The MV4-11 cells were transduced with 10µl of viral particles in a 96-well plate in the presence of 8µg/ml polybrene and centrifuged at 1600 rpm for 60 mins at room temperature on the first and second day. The transduced MV4-11 cells were transferred into fresh IMDM media at 48h and allowed to expand for another 1-2 days. The transduced mCherry-positive cells were sorted on a BD FACSAria III (BD Biosciences) flow-cytometry sorter and used for *in-vitro* luciferase expression validation and *in-vivo* transplantation experiments.

### Bioluminescence imaging (BLI) of Leukemia cells in vivo

For non-invasive assessment of leukemic burden, whole body imaging was performed every week using the LagoX Imaging System (Spectral Imaging, Tucson, AZ). Mice were injected with D-Luciferin solution (i.p. 150mg/kg), and 5 min later the mice were anesthetized with isoflurane (Faulding Pharmaceuticals) and imaged. Supine, prone and side view images were acquired for 10-30 sec and using AMIView software the photon emission transmitted from mice was captured and quantitated in photons/sec/cm$^2$/sr. Further analysis of the images was done using Aura 2.0.1 software.

### Blood clearance activity

For pharmacokinetic analysis of $^{111}$In-111 DOTA-anti-human CD33 antibodies, blood activities were converted to %ID/g and then normalized to the initial blood sample activity. Data from each time point was averaged together and a bi-exponential equation was fit to the data using nonlinear least-squares fitting in Matlab (version R2018a 9.4.0.813654, Mathworks, Natick MA) [2].

### Diagnostic accuracy

Sensitivity was calculated by taking the ratio of CD33+ AML mice that had a percent ID/g above a given threshold, to the total number of CD33+ mice. Specificity was calculated by taking the ratio of CD33- mice that had a percent ID/g below the same threshold to the total number of CD33- mice. The diagnosis percent ID/g threshold was set at 2.75 % for biodistribution data of whole left femur. All Mice in AML group (both MV4-11 and HL-60) were considered positive for CD33+ leukemia if they contained CD33+ cells in the left femoral bone marrow as determined by flow cytometry.

### References:

**1. Mallampati S, Sun B, Lu Y, Ma H, Gong Y, Wang D, et al. Integrated genetic approaches identify the molecular mechanisms of Sox4 in early B-cell development: intricate roles for RAG1/2 and CK1epsilon. Blood. 2014; 123: 4064-76.**

**2. Wang Z, Mårtensson L, Nilsson R, Bendahl P-O, Lindgren L, Ohlsson T, et al. Blood Pharmacokinetics of Various Monoclonal Antibodies Labeled with a New Trifunctional Chelating Reagent for Simultaneous Conjugation with 1,4,7,10-Tetraazacyclododecane-<em>N,N</em>′,<em>N</em>″,<em>N</em>‴-Tetraacetic Acid and Biotin before Radiolabeling. Clinical Cancer Research. 2005; 11: 7171s-7s.**

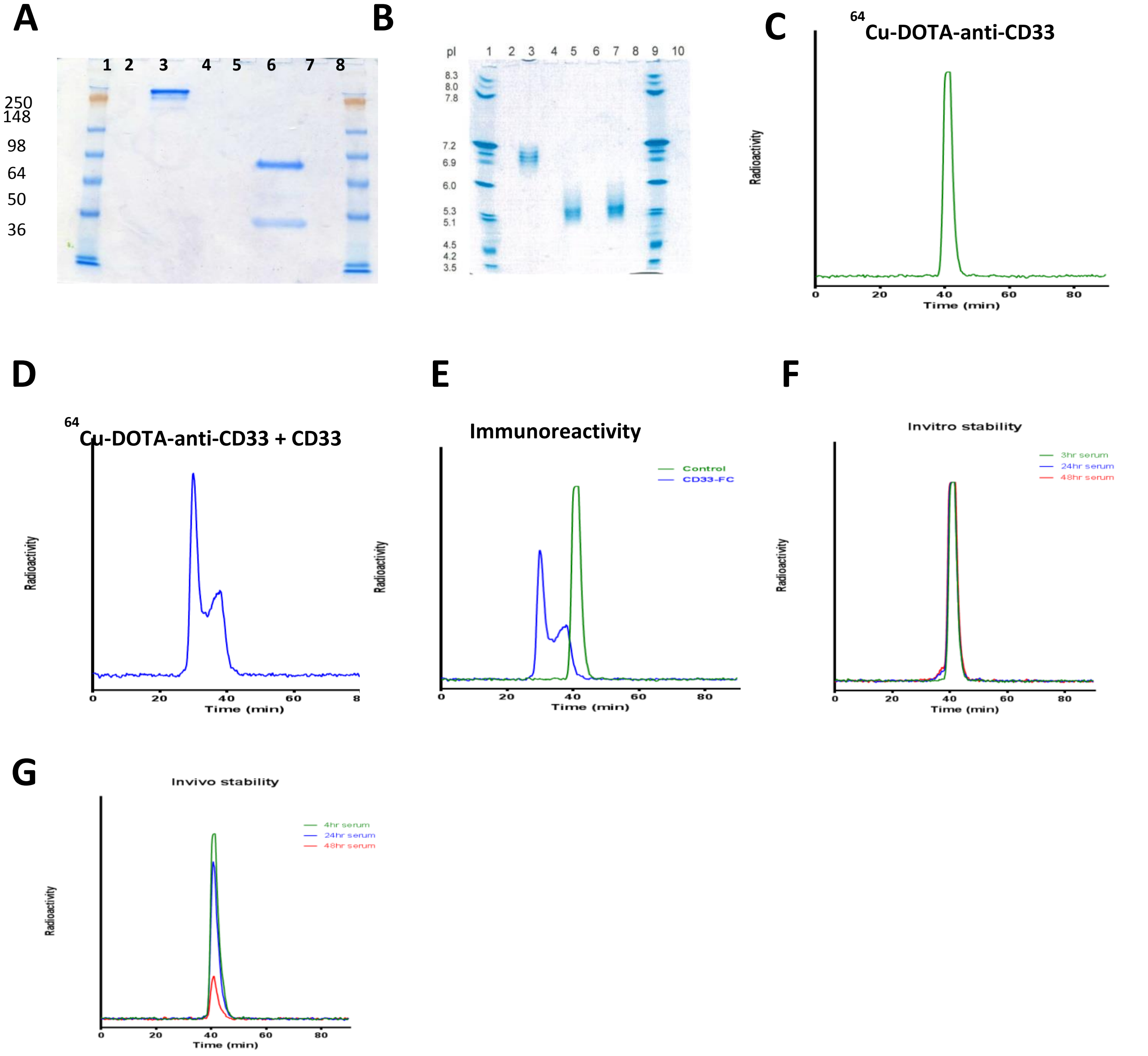


**Supplemental Figure S1**: **Anti-CD33 mAb antibody conjugated with DOTA, radiolabeling, immunoreactivity and stability.** **A**) The DOTA-anti-human CD33 mAb was electrophoresed on a SDS-PAGE gel under non-reducing (lane 3) and reducing conditions (lane 6) demonstrating purity (Lane 1 and 8, See blue plus 2 standard). **B)** The anti-human CD33 mAb (lane 3), DOTA-anti-human CD33 mAb (lane 5) and DOTA-anti-human CD33 mAb vialed product (lane 7) were analyzed on an iso-electrofocusing gel (Lane 1 and 9, Invitrogen IEF standards). Coomasie Blue staining showed a shift to a more acidic pH, confirming the conjugation process. The unconjugated anti-CD33 antibody showed a family of bands with an isoelectric point (pI) of > 6.9. Post-DOTA conjugation, the pI of anti-CD33-DOTA shifted to a more acidic range of ~5.3-5.6; **C)** Analysis of $^{64}$Cu-DOTA-anti-CD33 by HPLC size exclusion chromatography (SEC). Radiochromatogram (green) of the purified $^{64}$Cu-DOTA-anti-CD33 shows efficient labeling of anti-CD33-DOTA with Cu-64, with no aggregates and retention time around 40 min corresponding to an intact mAb. **D)** $^{64}$Cu-DOTA-anti-CD33 immunoreactivity. The purified $^{64}$Cu-DOTA-anti-CD33 was incubated with soluble CD33-Fc antigen and analyzed by SEC. The radioactivity peak (blue) showed a faster retention time ( ~30 min) indicating a shift to a higher molecular size consistent with binding to CD33 Fc antibody (67-85kDa). (**E**) Overlay of SEC radiochromatogram depicting a clear shift in $^{64}$Cu-DOTA-anti-CD33 incubated with CD33-Fc implying CD33 specific immunoreactivity. **F and G**) $^{64}$Cu-DOTA-anti-CD33 stability was tested *in vitro* and *in vivo* in mouse serum at different time points. SEC chromatogram clearly indicates the radiolabeled antibody was very stable even at 48h.

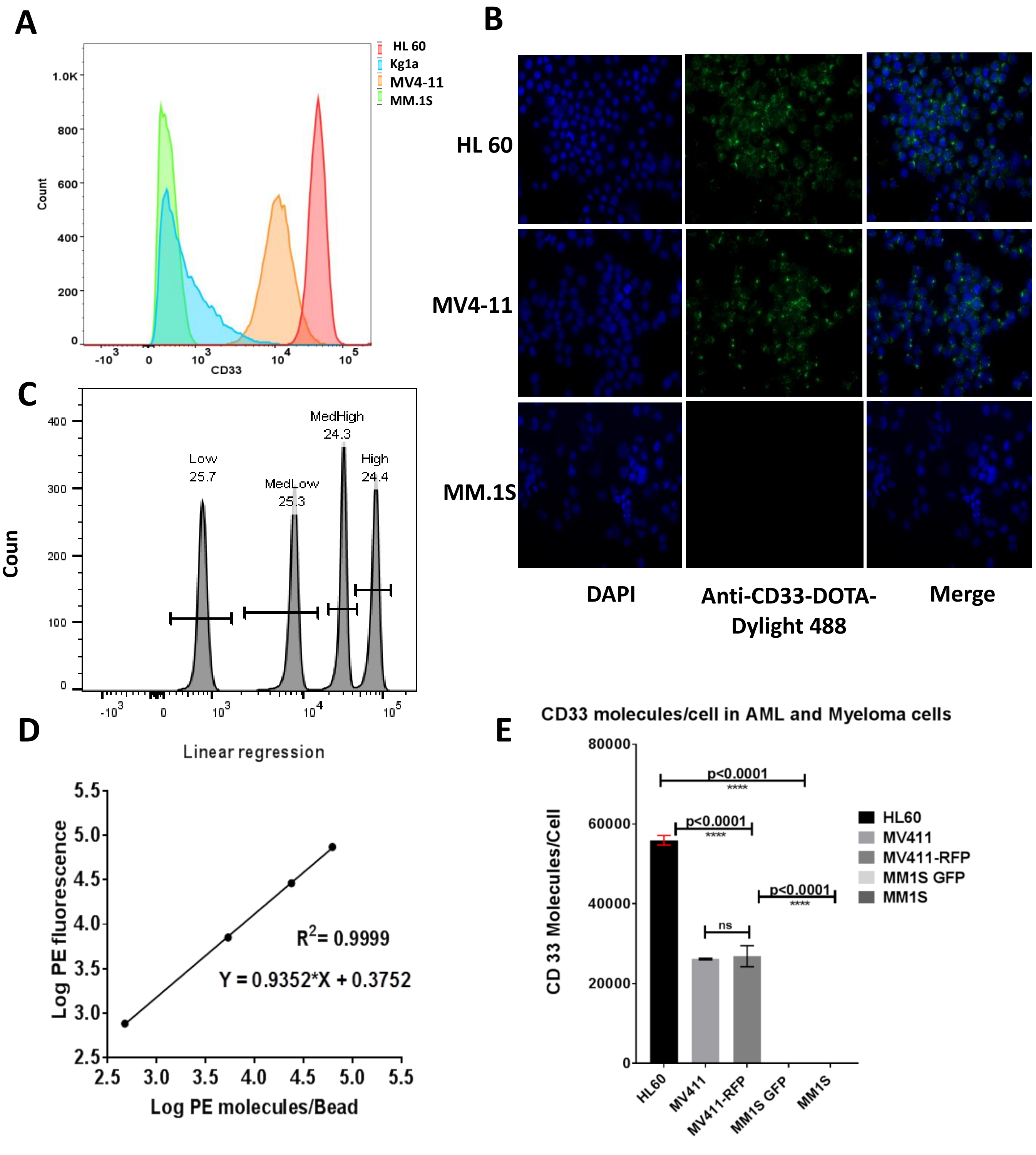


**Supplemental Figure S2: The anti-CD33-DOTA conjugated antibody immunoreactivity and quantification of CD33 antibody molecules per cell using BD QuantiBRITE PE.** (A) CD33 cell surface expression in AML and MM cell line using anti-CD33-DOTA dylight-488. MV4-11 and HL-60 cells were 100% positive for CD33 whereas Kg1a had <30% cells positive for CD33. Therefore, for further studies we only used MV4-11 and HL-60 AML cells. (B) CD33 immunofluorescence using anti-CD33-DOTA-dylight 488. AML and MM cells were stained with anti-CD33-DOTA-dylight 488. HL60 and MV4-11 showed CD33 immunofluorescence whereas negative control MM.1S had no CD33 staining. Unstained cells were used to set background, and all images were obtained with the same settings in Zeiss AxioObserver Z1 florescent microscope. (C) Histogram representing PE associated florescence (log values) and the interval gates were adjusted around each four bead peaks and labeled as Low, Med Low, Med High and High. (D) Linear regression plots for the number of PE molecules per bead (x axis) against fluorescence (y axis) (log10 values). (E) CD33 antibody per cell in AML and MM cell line. AML HL60 has more CD33 on cell surface than MV4-11 while MM.1S had no CD33 molecules.

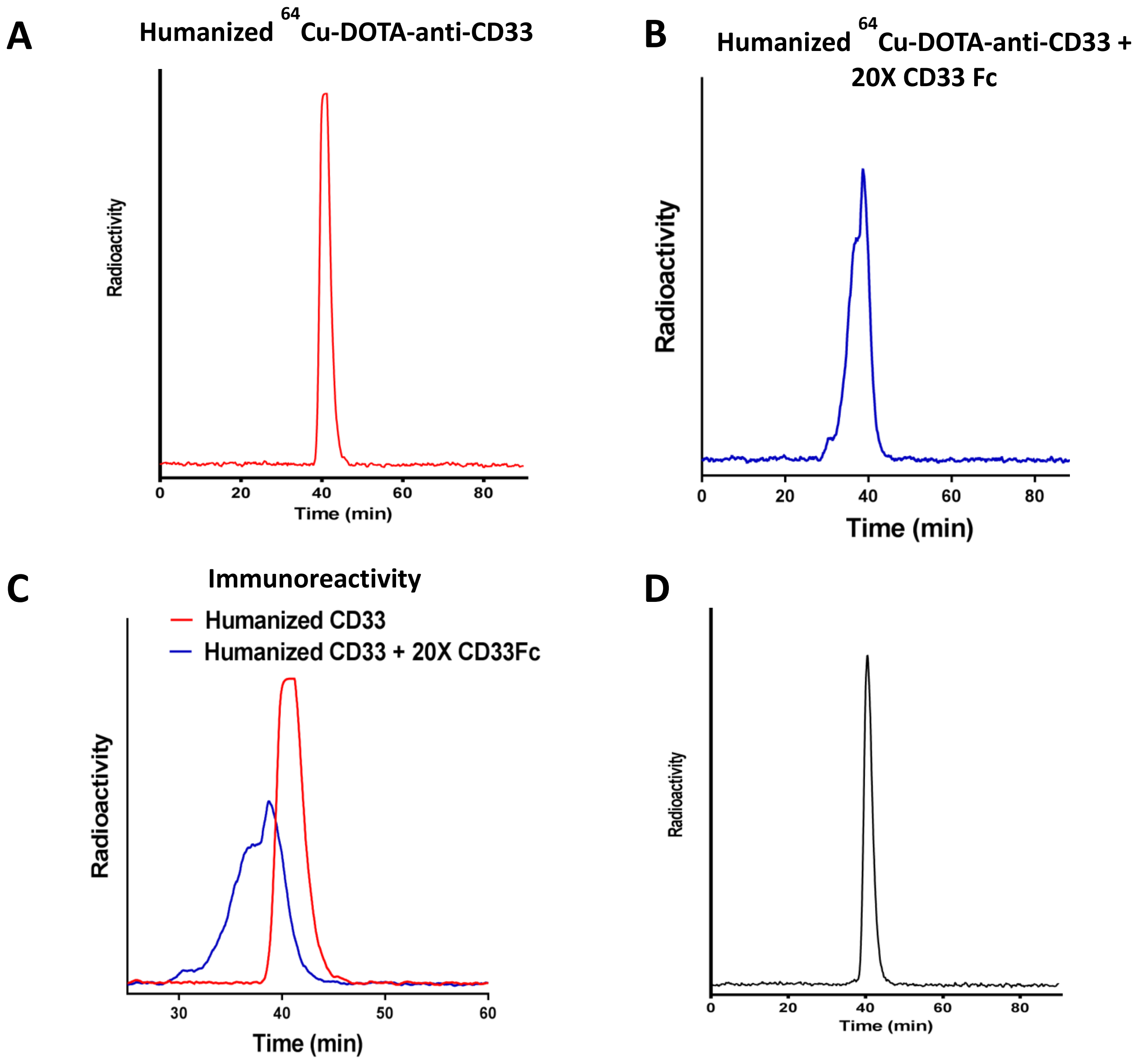


**Supplemental Figure S3**: **Humanized** DOTA-a**nti-CD33 mAb antibody radiolabeling, immunoreactivity and stability.** **A**) Analysis of humanized $^{64}$Cu-DOTA-anti-CD33 by HPLC size exclusion chromatography (SEC). Radiochromatogram (Red) of the purified $^{64}$Cu-DOTA-anti-CD33 shows efficient labeling of anti-CD33-DOTA with Cu-64, with no aggregates and a retention time around 40 min corresponding to an intact mAb. **B)** $^{64}$Cu-DOTA-anti-CD33 immunoreactivity. The purified $^{64}$Cu-DOTA-anti-CD33 was incubated with soluble CD33-Fc antigen and analyzed by SEC. The radioactivity peak (blue) showed a faster retention time (~35 min) indicating a shift to a higher molecular size consistent with binding to CD33 Fc antibody. **C**) Overlay of SEC radiochromatogram depicting a shift in $^{64}$Cu-DOTA-anti-CD33 incubated with CD33-Fc implying CD33 specific immunoreactivity. **D**) $^{64}$Cu-DOTA-anti-CD33 stability was tested *in vivo* in mouse serum at 48h and SEC chromatogram clearly indicates the radiolabeled antibody was very stable at 48 h.

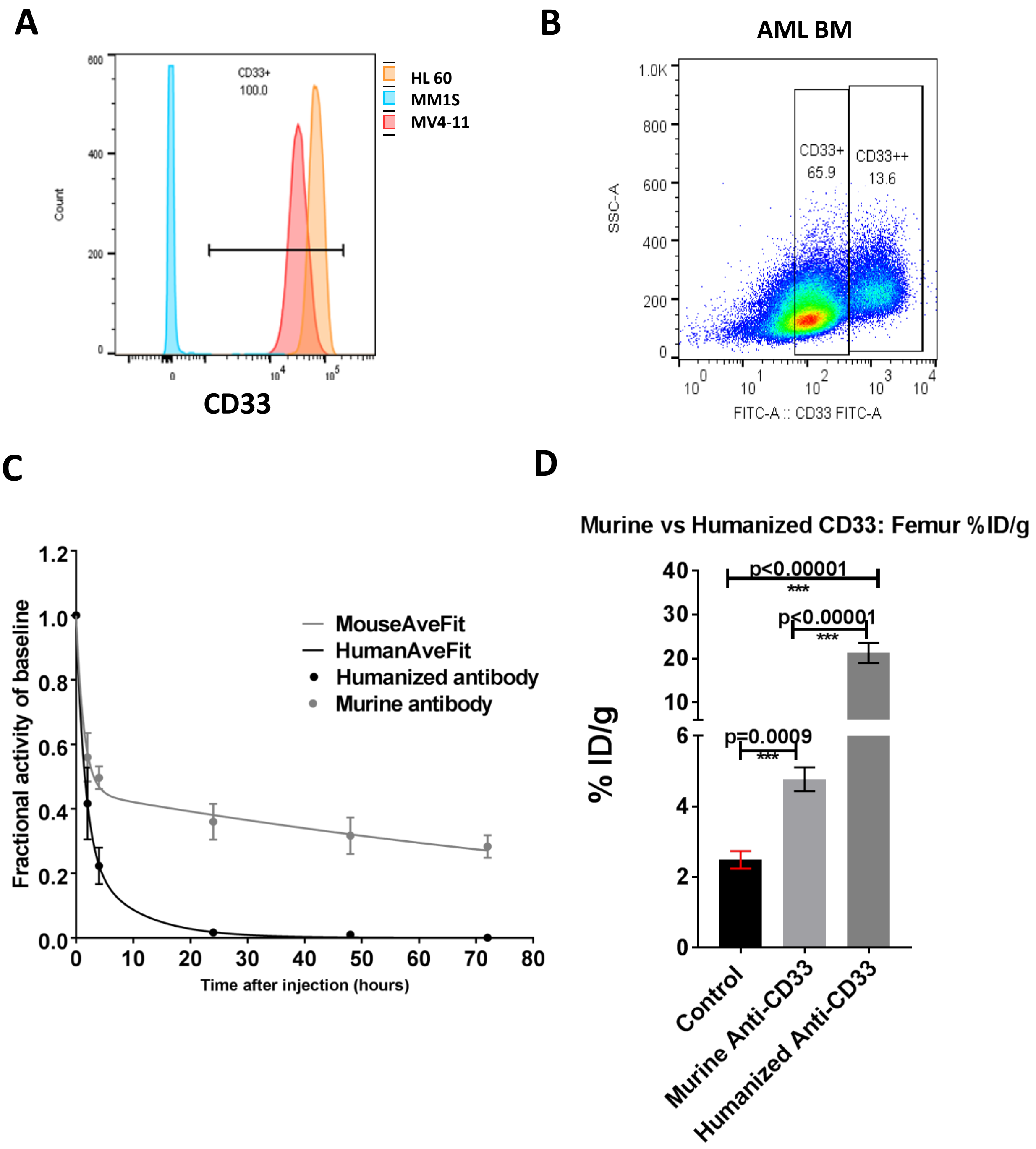


**Supplemental figure S4: Humanized anti-CD33 Immunoreactivity and blood activity clearance. A and B**) Humanized anti-CD33 immunoreactivity was determined in AML cell line and patient sample. The humanized anti-CD33 antibody detected CD33 in AML cells, human AML patients but not in CD33- MM.1S cells. **C)** Blood clearance activity of humanized and murine $^{64}$Cu-anti-CD33-DOTA anti-CD33 antibody was determined. Humanized mAb cleared faster with <5% activity in blood by 24h, while murine antibody maintained ~25% activity in blood for over 72h. The blood clearance activity was determined using dual exponential decay curve. **D)** The humanized Ab had better targeting to CD33+ AML than murine Ab, as shown by ~ 3-4 fold higher %ID/g (CD33 PET activity) in femur than murine Ab.

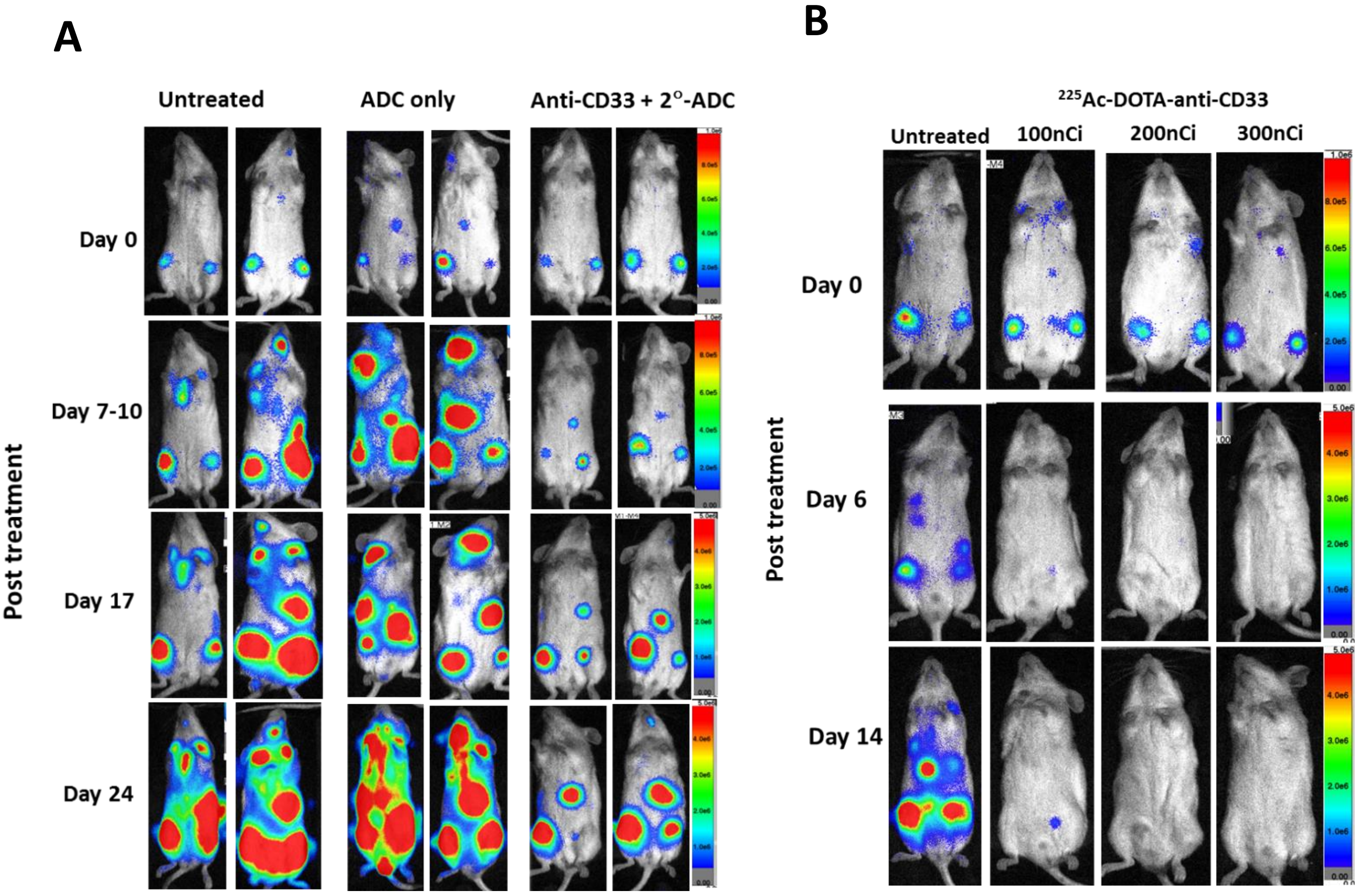


**Supplemental figure S5: Humanized anti-CD33 Therapeutic potential evaluated as an ADC and Radioimmunotherapy.** Therapeutic potential of humanized anti-CD33 mAb was evaluated as an ADC using anti-human Fc secondary Ab conjugated to MMAE. **A**) Representative BLI images of untreated, ADC alone treated and anti-CD33 + 2°ADC treated mice are shown. Anti-CD33 + 2°ADC treated mice clearly shows reduction in leukemia burden compared to respective controls. **B)** Radioimmunotherapy (RIT) using $^{225}$Ac-DOTA-anti-CD33 (CD33-RIT) was carried out in mice bearing MV4-11 AML. Representative BLI images of mice receiving $^{225}$Ac-DOTA-anti-CD33 at three different activities 100 nCi, 200 nCi and 300 nCi are shown. The disease burden is significantly reduced in all groups receiving CD33-RIT in comparison to untreated mice. The color scale bar represents the BLI intensity in photons/second/cm 2 /steradian (photons/s/cm$^2$/sr).
